# Genetic characterization of invasive house mouse populations on small islands

**DOI:** 10.1101/332064

**Authors:** Andrew P Morgan, John P Didion, Jonathan J Hughes, Jeremy B Searle, Wesley J Jolley, Karl J Campbell, David W Threadgill, Fernando Pardo-Manuel de Villena

## Abstract

House mice (*Mus musculus*) have dispersed to nearly every major landmass around the globe as a result of human activity. They are a highly successful invasive species, but their presence can be devastating for native ecosystems. This is particularly true on small offshore islands where mouse populations may grow unchecked by predators. Here we use genome-wide SNP genotypes to examine ancestry and population structure on two islands of ecological interest - Southeast Farallon Island, near San Francisco, CA; and Floreana Island in the Galápagos - in the context of a total cohort of 520 mice with diverse geographic origins, as a first step towards genetically-based eradication campaigns. We show that Farallon and Floreana mice, like those from previously-studied islands in both the Atlantic and Pacific Oceans, are of admixed European ancestry. We find that these populations are on average more inbred than mainland ones and passed through a strong colonization bottleneck with little subsequent genetic exchange. Finally we show that rodenticide resistance alleles present in parts of Europe are absent from all island populations studied. Our results add nuance to previous studies of island populations based on mitochondrial sequences or small numbers of microsatellites and will be useful for future eradication and monitoring efforts.

## Introduction

House mice (*Mus musculus*) are extremely successful as an invasive species. Due in large part to their commensal relationship with humans, house mice have dispersed from their ancestral range in central Eurasia to nearly every landmass visited by humans (reviewed in Bishop et al. (1985)). As frequent stowaways in shipside food stores, house mice have colonized numerous remote islands in both Atlantic and Pacific oceans, rivaling humans for global distribution. Their omnivorous and opportunistic feeding habits and short generation time allow them to quickly become established in a wide variety of environments (Berry 1970). On islands where they are largely free of natural predators, mice may develop a suite of life-history traits referred to as the “island syndrome“: increased body size, increased survival rates, decreased aggression and decreased dispersal (Adler and Levins 1994).

However, the arrival of house mice may be devastating for native ecosystems. They are considered among the most damaging invasive species (IUCN Global Invasive Species Database, http://www.iucngisd.org/gisd/100_worst.php). The best-studied impacts are on birds that rely on offshore islands as breeding grounds. In this work we focus on mice from two islands: Southeast Farallon Island, 45 km off the coast of San Francisco, CA; and Floreana Island in the Galápagos, Ecuador. The first documented landing on the Farallon Islands was by Sir Francis Drake in 1579, though the islands were named by the Spanish explorer Sebastián Vizcaíno in 1603 (White 1995). Russian and native Alaskan hunters periodically exploited seal populations throughout the 1800s, and egg-harvesting crews from San Francisco competed over collection territories in a bizarre episode known as the “Egg Wars” in the mid-1860s (Thompson 1896; White 1995). The islands (of which Southeast Farallon Island is the largest) were declared a National Wildlife Refuge in 1969 and are now off-limits to the public. They support the largest colony of breeding seabirds in the continental United States south of Alaska - more than 200,000 individuals representing at least 13 species in peak season - as well as an endemic species of cricket (Steiner 1989) and several species of seals, sea lions and sea otters. The mouse population on the Farallones is highly cyclical: breeding begins in late spring, populations peak in early autumn, and die-back occurs in winter as food becomes scarce. Peak densities exceed 1250 individuals per hectare (San Francisco Bay National Wildlife Refuge Complex 2013), at least an order of magnitude greater than the peak density of populations at nearby mainland sites (DeLong 1967). This abundance attracts raptor species, notably burrowing owls, that also prey on smaller birds (Mills 2016). Much less is known about the biology of house mice in the Galápagos, but it is likely that the presence of mice there has deleterious effects on native birds, plants and endemic rodents (Harper and Carrion 2011). Such threats to native ecosystems have motivated efforts to eradicate these invasive mammals. Although many mammalian eradications have been successful, more than 20% of attempted house mouse eradications using rodenticides have failed, and many more are not feasible due to the complexity of island ecosystems (Howald et al. 2007). This has lead to proposals for genetically-based approaches for eradication (Leitschuh et al. 2018).

Genetic characterization of island populations is a prerequisite to genetically-based eradication campaigns. Little is known about the origins, ancestry, or population structure of mice on either the Farallones or Floreana. Studies of several other offshore islands - Gough Island (Gray et al. 2014), the Faroe Islands (Jones et al. 2011), the Azores (Gabriel et al. 2015), the Kerguelen Archipelago (Hardouin et al. 2010), Antipodes (Searle et al. 2009; Veale et al. 2018), and Madeira (Gündüz et al. 2001) - have revealed predominantly western European ancestry consistent with human-mediated dispersal by mariners within the past several centuries. Those studies relied heavily on mitochondrial sequences with or without a small number of nuclear microsatellite markers, and thus had limited resolution for population structure and inbreeding.

Here we use genome-wide SNP data to explore the ancestry and population structure of mice from six groups of offshore islands, including Farallon and Floreana, and place them in the context of a larger cohort totaling 527 mice from across the globe. We confirm that mice from all islands sampled are overwhelmingly of western European *M. m. domesticus* origin. Geographic discordance between Y chromosome and mitochondrial markers on some islands provides evidence for inter-and intra-subspecific admixture that may have occurred before or after island colonization. We show that island mice are on average more inbred than their mainland counterparts, though the level of inbreeding varies greatly across mainland trapping sites. We use decay of linkage disequilibrium as a proxy for effective population size and show that different islands vary widely in the strength of the colonization bottleneck, the degree of ongoing genetic exchange with mainland populations, or both. Finally, we show that rodenticide-resistance alleles common in some European populations are absent from all islands examined, an important consideration for future eradication campaigns.

## Materials and methods

### Sample collection

Mice were collected at a large number of sites around the world (recorded in **Table S1**). All trapping and euthanization was conducted according to the animal use guidelines of the institutions with which providers were affiliated at the time of collection. Trapping on Southeast Farallon Island was carried out during two seasons, in 2011 and 2012; and on Floreana during a single season in 2012.

Samples of one or more tissues (including tail, liver, spleen, muscle and brain) from each individual were shipped to the University of North Carolina during a six-year period (2010-2016). We assigned each individual a unique identifier follows: `CC:LLL_SSSS:DD_UUUU` (`C` = ISO 3166-1-alpha2 country code; `L` = locality designation; `S` = nominal subspecies; `D` = diploid chromosome number and `U` = sequential numeric ID).

Genotypes for all individuals in this study have been used in prior studies on a selfish genetic element in European *M. m. domesticus* (Didion et al. 2016) and/or a *M. m. domesticus* - *M. m. castaneus* hybrid zone in New Zealand (Veale et al. 2018).

### DNA preparation and genotyping

Whole-genomic DNA was isolated from tissue samples using Qiagen Gentra Puregene or DNeasy Blood & Tissue kits according to the manufacturer’s instructions. Genotyping was performed using either the Mega Mouse Universal Genotyping Array (MegaMUGA) or its successor, GigaMUGA (GeneSeek, Lincoln, NE). Genotypes were called using Illumina BeadStudio (Illumina Inc, Carlsbad, CA). Only samples with < 10% missing calls were retained for analysis.

Sample sexes were confirmed by comparing the number of non-missing calls on the Y chromosome to the number of heterozygous calls on the X chromosome, which we have previously shown to be an effective means of discriminating genetically male from female samples.

### Genotype calling from whole-genome sequencing

Whole-genome sequence (WGS) data was obtained from the European Nucleotide Archive for 67 mice representing natural populations of *M. m. domesticus* (PRJEB9450), *M. m. castaneus* (PRJEB2176), *M. m. musculus* (PRJEB14167, PRJEB11742) and *M. spretus* (PRJEB11742) (Harr et al. 2016), plus one *M. spicilegus* (PRJEB11513) (Neme and Tautz 2016) and one *M. caroli* individual (PRJEB14895) as outgroups. All mice were either directly trapped in the wild or were first-generation laboratory offspring of wild-caught parents. Raw reads were aligned to the mm10 mouse reference genome with `bwa mem` v0.7.15; optical duplicates were marked with `samblaster` v0.1.22 and excluded from further analyses. Genotypes were called with the Genome Analysis Toolkit (`GATK` v3.3-0-g37228af) `UnifiedGenotyper` (McKenna et al. 2010) in all 67 individuals jointly. We attempted to call only the 77 726 sites targeted by MegaMUGA whose probe sequences mapped to the reference genome (with `BLAT`, and excluding unplaced contigs in the reference), but allowed for the discovery of additional alternate alleles besides those targeted on the array.

### Genotype merging and filtering

Raw genotypes from the MegaMUGA (*n* = 310 individuals), GigaMUGA (*n* = 150), and WGS (*n* = 67) were merged in hierarchical fashion with the aim of minimizing systematic differences in call rates across platforms. The two SNP array sets were merged first, as the GigaMUGA platform is almost a superset of the most reliably-called markers on MegaMUGA (Morgan et al. 2016). First, any samples with ≥ 10% missing calls were dropped. Next all markers present on both MegaMUGA and GigaMUGA, reported on the same strand on both platforms, and with probe sequences mapping to the autosomes, X, Y or mitochondrial genomes (*m* = 66 397) were retained and all other markers were dropped. Alleles were updated to encode all genotypes with respect to the reference allele on the plus strand, dropping sites with potential ambiguity in strand or physical position (*m* = 58 772).

Ascertainment bias on the MegaMUGA and GigaMUGA arrays is severe, owing both to design goals and the idiosyncratic history of laboratory strains of mice. Most array content was ascertained in laboratory strains among which *M. m. domesticus* ancestry is heavily over-represented, and alleles tagged by the array and segregating in *M. m. musculus* or *M. m. castaneus* are likely to be older and ancestrally polymorphic. The sites targeted by the MegaMUGA and GigaMUGA arrays are heavily biased towards transitions (Ti:Tv = 4.1) and a large fraction (23%) are CpG sites. Copy-number variation or other off-target variants in or near the probe sequence interfere with hybridization and may create systematic error in genotype calls (Didion et al. 2012). The consequences of these biases for population genetic analysis are discussed in detail in a companion manuscript (Hughes, Morgan, Didion *et al.*, in preparation).

Briefly, we used WGS to discover and mitigate possible biases due to recurrent mutation or copy-number variation. Sites with multiple alternate alleles among the 67 WGS samples (which include representatives of outrgroups *M. spretus*, *M. spicilegus* and *M. caroli*) were flagged as potential recurrent mutation or low-quality calls (4 374). Ancestral alleles were inferred as the majority-rule consensus call in *M. caroli* and *M. spicilegus* and sites where the derived allele is present in both *M. musculus* and either *M. spicilegus* and *M. caroli* were dropped as potential recurrent mutations (3724). WGS and array datasets were then merged. Finally, 600 sites with systematic difference in call rate (Fisher’s exact *p*, 0.001) were dropped. These filtering steps yielded a genotype matrix of 52 280 sites × 520 samples with final genotyping rate 97.5%.

The merged genotype matrix is available as **File S1** in VCF format.

### Relatedness

Cryptic relatives were identified on the basis of autosomal genotypes with akt kin v3beb346 (Arthur et al. 2017) using the “KING-robust” kinship estimator (Manichaikul et al. 2010). Kinship estimation was performed separately in each taxon (*M. m. domesticus*, *M. m. musculus*, *M. m. castaneus*, *M. m. gentilulus*, *M. spretus*) because allele frequencies are highly stratified between these groups. For each pair with kinship coefficient > 0.10, one member was removed at random to yield a set of 403 putatively-unrelated individuals. The genotype matrix including only unrelated individuals is available as **File S2** in VCF format.

### Analyses of population structure

Mitochondrial (M) and Y chromosome haplogroups were assigned using a clustering approach. Heterozygous genotypes were marked missing since both Y and M are hemizygous. Markers with > 7.5% missing data were dropped, leaving 13 markers on Y and 16 markers on M. Principal component analysis (PCA) was performed on hemizygous genotypes for all individuals (M) or all males (Y) using the `R` package `argyle` v0.2 (Morgan 2016). The number of clusters was determined by manual inspection. (For Y, four males with an excess of missing calls on Y were identified as outliers in the initial PCA and removed the analysis.) Individuals were then assigned to clusters using the partitioning-around-medioids (PAM) algorithm as implemented in `R` package `cluster` v2.0.5 (https://cran.r-project.org/web/packages/cluster/).

PCA on autosomal genotypes was performed using the randomized SVD algorithm implemented in `akt pca` v3beb346 (Arthur et al. 2017) and computing only the top 20 PCs. To avoid distortion of genotype space by groups of related individuals, PCs and loadings were calculated over the set of unrelated individuals only; remaining samples were then projected onto these coordinates.

Subspecies-level ancestry proportions were estimated using `ADMIXTURE` v1.3 (Alexander et al. 2009) in supervised mode with *K* = 3 components. We first defined a set of 9 273 ancestry-informative markers (8 875 autosomal, 401 X-linked) for which the derived allele is present at moderate frequency in a single subspecies, as described elsewhere (Hughes, Morgan, Didion *et al.*, submitted). A subset of 73 unrelated individuals (43 *M. m. domesticus*, 14 *M. m. musculus*, 16 *M. m. castaneus*) known on the basis of prior studies to have pure subspecies ancestry were used as the training set, and the remaining unrelated individuals as unknowns. Standard errors on ancestry proportions were calculated using `ADMIXTURE`’s bootstrap procedure with 100 rounds of resampling.

Further ancestry analyses within *M. m. domesticus* were performed by running `ADMIXTURE` in unsupervised mode with *K* = 2,…, 12 components. Here we used a pruned genotype matrix containing variants at within-subspecies MAF > 1% (38 018 autosomal sites, 1001 X-linked sites) to improve stability of ancestry estimates.

The “population tree” in **Figure 3** was produced with `TreeMix` v1.12r231 (Pickrell and Pritchard 2012). Sites were first pruned for LD using `PLINK` v1.90p (command `––indep-pairwise 500 1 0.5`). Seven *M.spretus* individuals from Spain were used as outgroup to root trees. Runs were performed with *m* = 0, …, 5 gene-flow edges. Outgroup *f*_3_-statistics in **Figure 3** were calculated with the `threepop` utility and four-population *f*_4_-statistics for admixture graphs with the `fourpop` utility, both included in the `TreeMix` suite; standard errors were calculated by block jackknife over blocks of 500 sites.

We fit admixture graphs in an effort to understand the relationship of the Farallon and Floreana populations to populations representative of southern Europe (Spain), the Mediterranean (Greece), northern Europe (Maryland) and an outgroup (Iran) (**Figure S1**). Graph topologies were specified manually and edge weights were fit to *f*_4_-statistics (ie. *D*-statistics) using the maximum-likelihood procedure implemented in the `fit_graph()` function in `R` package `admixturegraph` v1.0.2 (Leppälä et al. 2017). Graphs were scored on the basis of the sum of squared residuals between the fitted edge weights and those implied by the observed /4-statistics. Axmixture edges were said to be estimable if constraining them to their fitted value did not reduce the sum of squared errors.

### Identity-by-descent

Genomic segments putatively shared identical-by-descent (IBD) between individuals were identified with the `fastIBD` module of `BEAGLE` v4.1-r7e1 (Browning and Browning 2013) with default parameters except for setting `ibdtrim=2` (number of markers arbitrarily clipped from ends of putative IBD segments). The resulting segments were then polished with the `merge-ibd-segments` v249 utility included with `BEAGLE`, merging segments < 0.01 cM apart and with at most 2 discordant genotype calls in the merged segments. The analysis included the full set of 520 samples without filtering for relatedness, as pairs of close relatives provide useful “hints” to the phasing algorithm. We were unable to run `BEAGLE` in sex-aware mode, so IBD segments are reported for autosomes only.

### Inbreeding

Individual inbreeding coefficients (*F̂*) were estimated as the fraction of the autosomes covered by runs of homozygosity (ROH). This has previously been shown to be a reasonably consistent estimator of recent inbreed-ing (McQuillan et al. 2008). ROH were identified separately in each individual using `bcftools roh` v1.7 (Narasimhan et al. 2016) with taxon-specific allele frequencies (estimated over unrelated individuals only in each of *M. m. domesticus*, *M. m. musculus*, *M. m. castaneus*, *M. m. gentilulus* and *M. spretus*), constant recombination rate 0.52 cM/Mb (Liu et al. 2014), and arbitrarily assigning array genotypes a Phred-scale GT = 30 (corresponding to error rate 0.001) per the authors’ recommendations. Other HMM parameters were left at their defaults after experimenting with a wide range of parameter settings, and finding the root mean squared error (relative to estimates of *F̂* from WGS data) to be insensitive to these parameters. Mean and median length of putative ROH segments were 10.0 Mb (6 098 sites) and 4.1 Mb (5 411 sites) respectively.

### Linkage disequilibrium

We estimated linkage disequilibrium (LD) between all pairs of autosomal markers on each chromosome, taking the mean value in 10 kb bins of inter-marker distance. LD calculations for each chromosome and population were performed with `PLINK` v1.90p (command `––make-founders ––ld-window-r2 0.0000000000001 ––r2 inter` Physical distance was scaled to genetic distance using recombination rate 0.52 cM/Mb. The equation of Sved (1971) was fit by non-linear least squares using the `nls()` function in base `R`, yielding a model-based estimate of *N*_*e*_.

Coalescent simulations were used to evaluate the effect of specific demographic histories on patterns of LD decay. These were implemented in Python using `msprime` (Kelleher et al. 2016). We simulated an island-mainland model in which an ancestral “mainland” population (*N*_*e*_ = 10000) gives rise to an “island” population with an associated bottleneck (reduction in *N*_*e*_ by 1/*f*) at *τ* generations in the past, with continuous symmetric migration at rate m between island and mainland. For each scenario we simulated 1 Mb of sequence with mutation rate 1 × 10^−8^ per bp per generation and recombination rate 0.5 cm/Mb. A sample of 20 chromosomes (10 diploid individuals) was drawn from the result, and binned LD calculated as from the real data.

### Rodenticide resistance

From five derived substitutions in *Vkorc1* associated with warfarin resistance in mice (Song et al. 2011), two are directly genotyped by the MegaMUGA and GigaMUGA arrays: chr7:127895440 C>T (Ensembl transcript ENSMUST00000033074.6: c.142G>A / p.Ala48Thr) and chr7:127894606 C>A (ENSMUST00000033074.6: c.182G>T / p.Arg61Leu). We estimated their frequency among unrelated individuals in the populations shown in **Figure 6** and calculated 95% highest posterior density intervals using the function `binom.bayes()` in `R` package `binom` v1.1-1 (https://cran.r-project.org/package=binom). The heatmap of frequencies for the allele p.Arg61Leu allele (**Figure 6**) was created by taking computing the centroid of each European population and then interpolating allele frequency in space via ordinary kriging on a spherical variogram, via functions `vgm(…, model=’Sph’)` and `krige()` in the `R` package `gstat` (Pebesma 2004).

## Results

Our dataset consists of 520 mice trapped in 32 countries between 1990 and 2015 (**Table 1** and **Table S1**). The collection spans the three principal subspecies of the house mouse across its native range (414 *M. m. domesticus*, 28 *M. m. musculus* and 39 *M. m. castaneus*) as well as the outgroup species M. *spretus* (8), M. *spicilegus* (1) and *M. caroli* (1), and a group with ill-defined ancestry provisionally labelled *M. m. gentilulus* (27) (**Figure 1**). The majority were genotyped with one of a pair of Illumina Infinium SNP arrays (Morgan et al. 2016) - the 78K-site MegaMUGA (310, 60%) or 143K-site GigaMUGA (143, 28%) - and the remainder were obtained from published whole-genome sequencing (WGS) data (67, 13%) (Harr et al. 2016). Briefly, the two array datasets were merged on overlapping markers, and WGS samples were added by calling genotypes at array sites. WGS data was also used to identify and remove problematic sites. Full details of data processing and quality control are provided in the **Materials and methods**. The final genotype matrix covers 52280 sites (50551 autosomal, 1689 X chromosome, 24 Y chromosome, 16 mitochondrial) with overall genotyping rate 97.5%.

**Table 1:** Counts of individuals used in this study, by taxon and population. Full sample descriptions provided in **Table S1**. For populations defined by an island, land area for that island is shown.

| Taxon | Population | males | females | N | unrelated | Land area (mi<sup>2</sup>) |
| --- | --- | --- | --- | --- | --- | --- |
| *M. m. domesticus* | Antipodes | 3 | 1 | 4 | 4 | 8.5 |
|  | Auckland | 3 | 1 | 4 | 4 | 197 |
|  | Australia | 10 | 0 | 10 | 5 |  |
|  | Belgium | 0 | 2 | 2 | 2 |  |
|  | Chatham | 2 | 2 | 4 | 3 | 355 |
|  | Cologne | 7 | 0 | 7 | 7 |  |
|  | Cyprus | 5 | 2 | 7 | 7 |  |
|  | Farallon | 13 | 7 | 20 | 17 | 0.3 |
|  | Floreana | 8 | 7 | 15 | 6 | 66.8 |
|  | France | 14 | 6 | 20 | 18 |  |
|  | Greece | 21 | 7 | 28 | 28 |  |
|  | Heligoland | 1 | 2 | 3 | 1 | 0.65 |
|  | Holstein | 3 | 0 | 3 | 3 |  |
|  | Iran | 8 | 0 | 8 | 5 |  |
|  | Italy | 2 | 4 | 6 | 6 |  |
|  | Lebanon | 1 | 3 | 4 | 4 |  |
|  | Madeira | 1 | 0 | 1 | 1 | 286 |
|  | Maryland | 49 | 62 | 111 | 50 |  |
|  | New Zealand North | 24 | 19 | 43 | 39 |  |
|  | New Zealand South | 36 | 35 | 71 | 56 |  |
|  | Orkney | 0 | 2 | 2 | 2 | †6.1 |

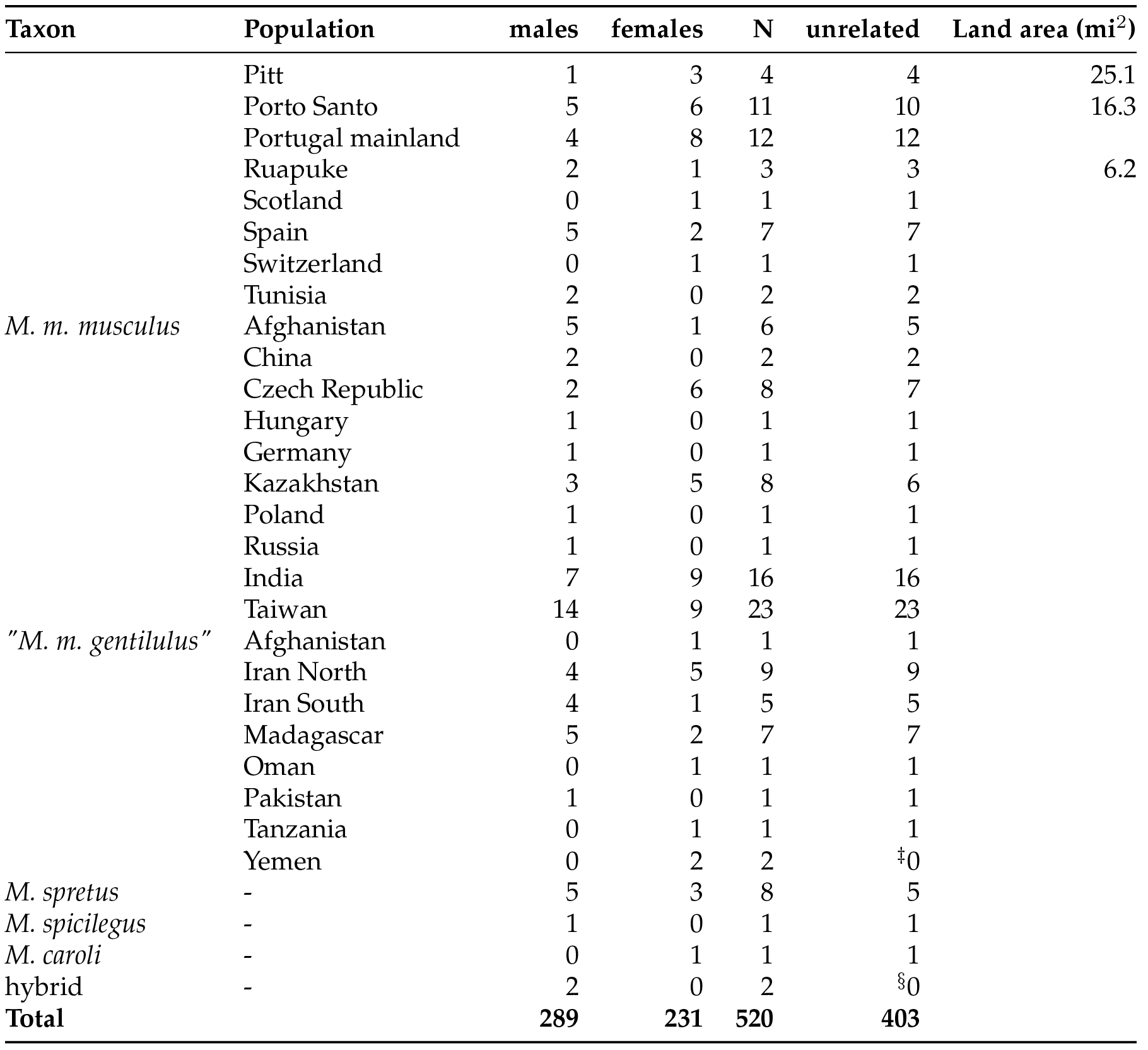

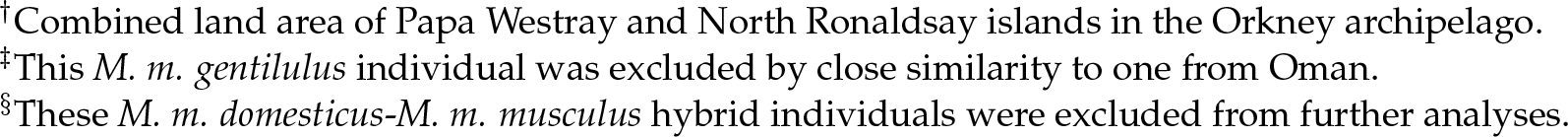

Many population-genetic analyses assume that individuals are not closely related. We identified and removed putative relatives on the basis of autosomal genotypes, leaving 403 putatively-unrelated individuals (173 female, 230 male).

Nine offshore islands, in six geographic groups, are represented: Southeast Farallon Island (17 unrelated mice); Floreana Island (6); Antipodes, Auckland, Chatham and Pitt Islands, in the Southern Ocean (15) (Veale et al. 2018); Heligoland, off the German coast (1); the Orkneys, north of Scotland (2); and Madeira and Porto Santo in the mid-Atlantic (11).

### Ancestry and population structure

We inspected global ancestry patterns by performing principal components analysis (PCA) on autosomal sites (**Figure 2A**). The top two principal components clearly partition individuals according to their nominal ancestry at the subspecies level. Supervised admixture analysis with three components trained on known pure representatives of each subspecies confirmed that all island populations have, on average, > 99% *M. m. domesticus* ancestry on both autosomes and the X-chromosome (**Table S2**). Two individuals, both from the Southern Ocean, have statistically significant *M. m. musculus* ancestry, but the absolute quantity is small (< 0.35%). No island mice have detectable *M. m. castaneus* ancestry for biparentally inherited markers. Mitochondrial and Y chromosome haplogroups are summarized in **Table S3**. As previously reported by Veale et al. (2018), *M. m. castaneus* mitochondrial haplogroups are widespread in New Zealand and are present in mice from Chatham Island, evidence of historical contact with southeast Asian populations. The *M. m. domesticus* mitochondrial haplogroup otherwise predominates on all islands examined. All island males carry *M. m. domesticus* Y chromosomes. We can further define three Y-chromosome haplogroups within *M. m. domesticus:* “Europe 1”, “Europe 2” and “Mediterranean.” Mice from Southeast Fara llon Islands carry “Mediterranean” exclusively; only “Europe 1” is present on Floreana. All three haplogroups are present in the Southern Ocean.

We next investigated the relationship between island populations and mainland groups within *M. m. domesticus* using autosomal markers. A heuristic population tree was constructed with TreeMix (Pickrell and Pritchard 2012) (**Figure 3A**). Putative relationships were quantitatively assessed with outgroup *f*_3_ statistics, which are relatively robust to SNP ascertainment (Peter 2016). As expected, Porto Santo mice have strong affinity to populations from Spain and Portugal. Mice from the islands of the Southern Ocean (**Figure 3B**, top row) have clear affinity for northern European, North American, and New Zealand populations (of probable British ancestry (Searle et al. 2009)), although the effect is less pronounced on Antipodes. The Farallon mice have greater affinity for northern European-derived populations than for Mediterranean or Iberian ones (**Figure 3B**, bottom row), but also have a distinct ancestry component not well-captured in our set of reference populations (see **Figure 2B,C**). Mice on Floreana have a similar profile to the Farallon mice, but with a larger Iberian component. We attempted to clarify the origins of Farallon and Floreana mice via model-based fitting of admixture graphs with representative reference populations (**Figure S1**). The best-fitting models place both the Farallon and Floreana populations as descendants of admixture events between the ancestors of present-day North American mice and the ancestors of present-day Spanish mice, but we were unable to obtain stable estimates of admixture proportions.

Global ancestry analyses using the grade-of-membership model implemented in ADMIXTURE (Alexander et al. 2009) largely recapitulate the patterns evident in PCA projections. For *K* = 2 (where *K* is the number of ancestry components in the model), we recover components that roughly correspond to “northern Europe” and “southern Europe/Mediterranean” (**Figure S2**). The “northern Europe” component is also evident in regions colonized by northern Europeans including the eastern United States and New Zealand. At *K* = 3 a component unique to the Southeast Farallon mice emerges; the Farallon mice remain distinct for all *K* ≥ 3 (**Figure S3**). By contrast, the Floreana mice have an ancestry profile intermediate between “northern European” populations (*eg.* Germany, New Zealand) and Iberian ones (Spain, mainland Portugal). With the exception of a single outlier individual trapped on Porto Santo (`PT:PSA_STND:40_0874`), island populations are genetically cohesive. Small sample sizes on each island limit the strength of the conclusions that can be drawn from this observation.

**Figure 1:**
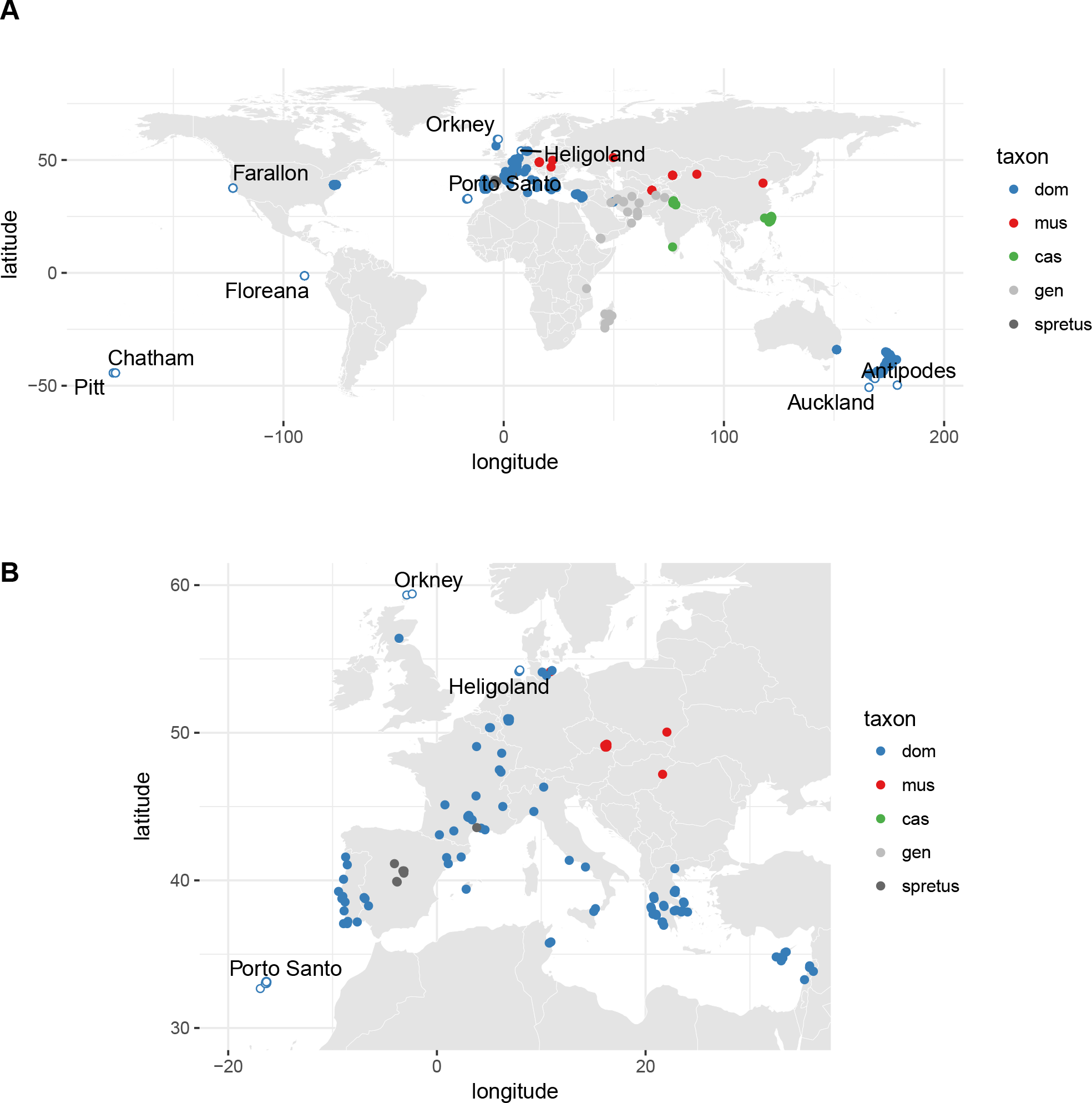
Geographic distribution of mice used in this study. **(A)** All samples, colored by nominal subspecies of origin (dom = *M. m. domesticus*, mus = *M. m. musculus*, cas = *M. m. castaneus, gen* = “*M. m. gentilulus”/uncertain*, spretus = *M. spretus.* Open dots, isolated island mice (labeled); closed dots, mainland mice. **(B)** Zoomed-in view of Europe and Mediterranean.

**Figure 2:**
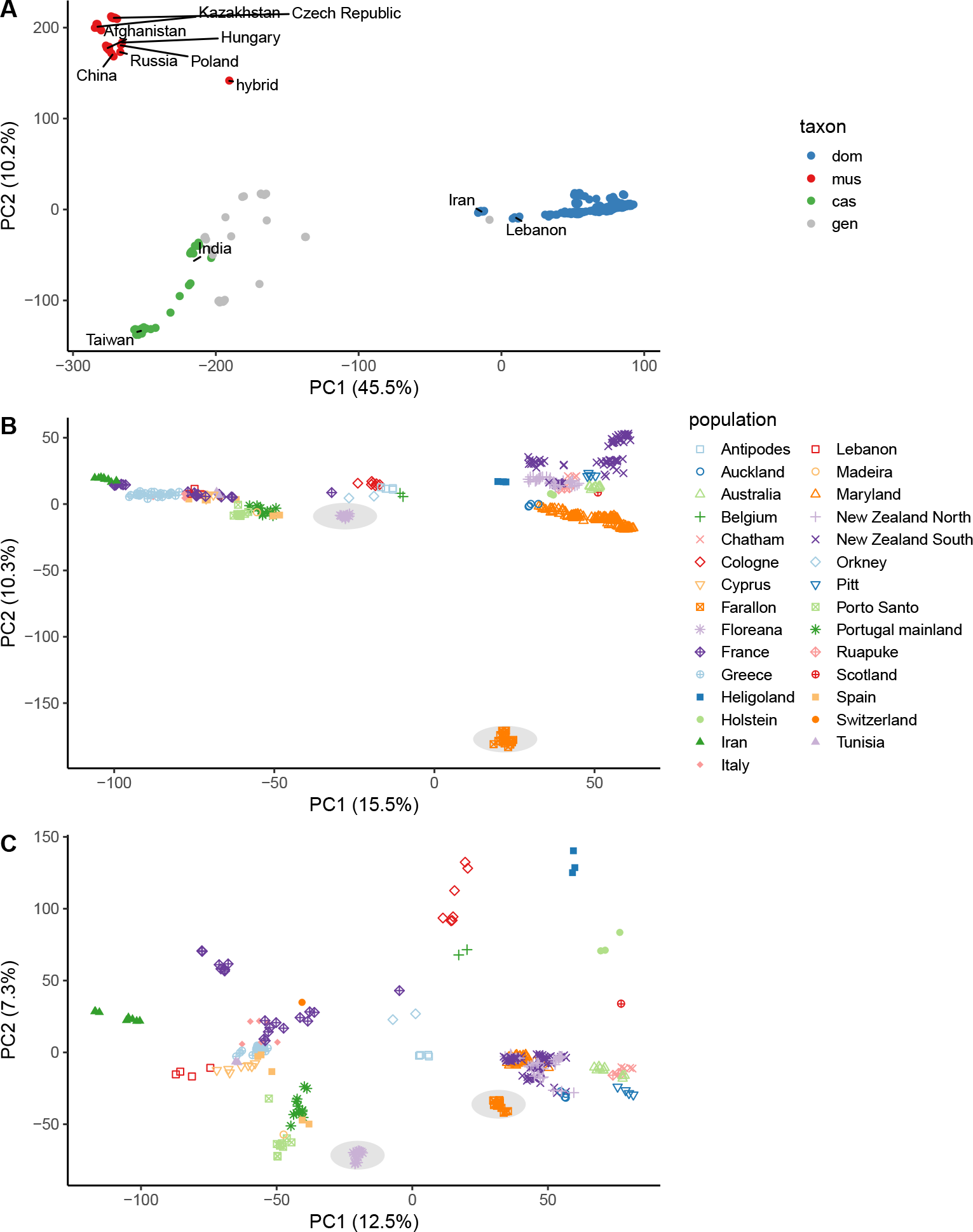
Principal components analysis (PCA) on autosomal genotypes. PCA was performed with unrelated samples only, and then all samples were projected onto the resulting eigenvectors. Proportion of variance explained by each axis is indicated in axis label. **(A)** All samples, colored by nominal taxonomic origin as in **Figure 1. (B)** *M. m. domesticus* samples only, with unique symbol-color combination per population. **(C)** *M. m. domesticus* samples only, with correction to account for unequal sampling effort across geographic locations. Symbols and colors as in panel B. Farallon and Floreana populations are emphasized with underlying grey shading in panels B and C.

**Figure 3:**
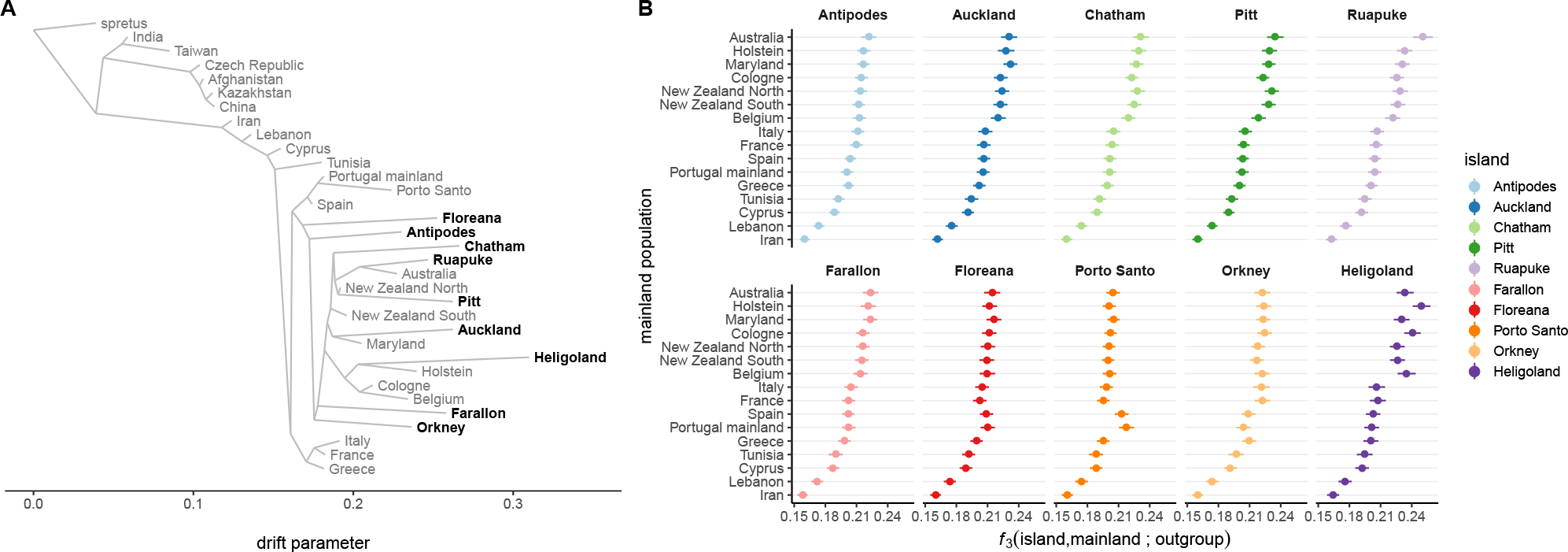
Patterns of admixture in mice from isolated islands. **(A)** Population tree for M. m. *domesticus* inferred by `TreeMix`. **(B)** Outgroup *f*_3_-statistics for island populations, measuring degree of shared genetic drift between each query population (one per panel) and each source population (on y-axis.)

As an alternative to models of population structure based on allele frequencies at unlinked sites, we also examined long haplotypes putatively shared identical by descent (IBD). We focused on IBD segments longer than 1.5 cM - that is, segments with expected time to coalescence of 33 generations. These segments provide a window onto shared ancestry at shallower time depths than allele-frequency differences (**Figure S4**). We find that mice from Floreana Island share recent co-ancestry with mice from Spain, mainland Portugal and Porto Santo․․ Floreana mice have no clear affinity to any mainland population on the basis of IBD segments. Both populations, however, have substantial recent co-ancestry with mice from the Southern Ocean. This pattern is consistent with wide dispersal of a northern European ancestry component by human activity within the past several hundred years.

**Figure 4:**
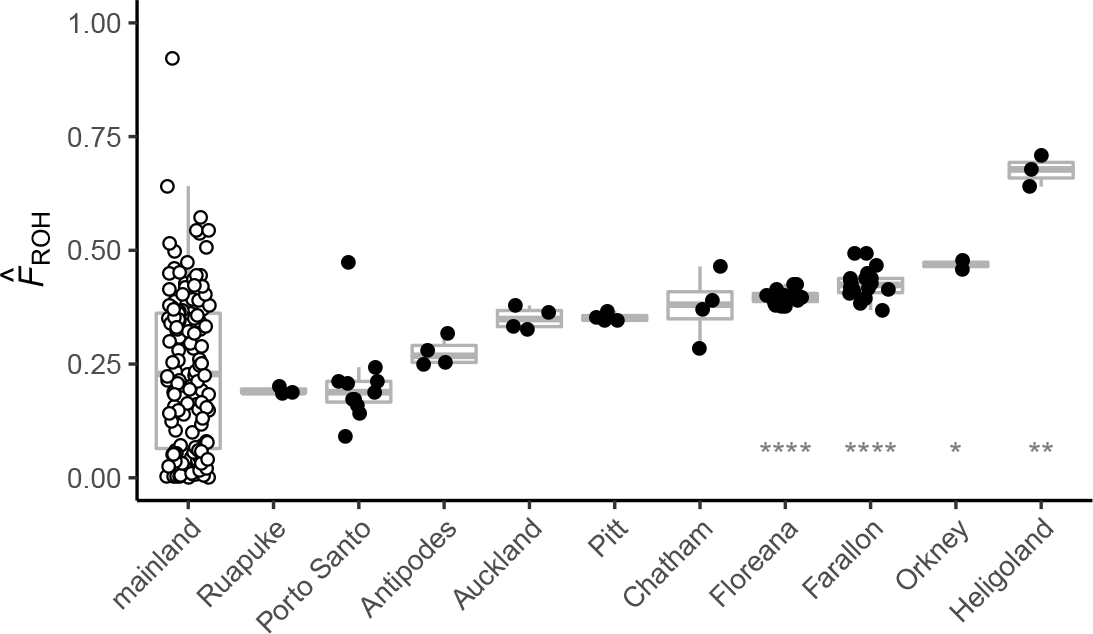
Individual inbreeding coefficients, estimated as the proportion of the genome contained in runs of homozygosity (*F̂*), in island populations (closed dots) versus mainland populations (open dots). Pairwise Wilcoxon rank-sum tests were performed between each island population and the mainland as a reference group; populations with significantly greater inbreeding than the mainland are marked above the axis (* = *p* < 0.05, ** = *p* < 0.01, *** = *p* < 0.001, **** = *p* < 0.0001).

### Inbreeding

Individual inbreeding coefficients were estimated by calculating the fraction of the autosomes contained in runs of homozygosity (ROH) with a model calibrated against subspecies-specific allele frequencies. House mouse populations vary widely in their level of inbreeding, from almost complete outcrossing to near homozygosity (**Figure 4**). Inbreeding is variable between islands (from *F̂* = 0.21 ± 0.10 [mean ± SD] on Porto Santo to *F̂* = 0.67±0.03 on Heligoland) but is generally consistent within them. Island mice are, as a group, significantly more inbred than mainland counterparts (*p* = 3.6 × 10^−9^, Wilcoxon rank-sum test). Mice from Floreana Island (*F̂* = 0.40±0.02) and Southeast Farallon Island (*F̂* = 0.43±0.03) rank among the most inbred populations in our study, significantly more so than mainland mice taken as a whole (*p* < 0.0001 for both comparisons, Wilcoxon rank-sum test). We might expect that inbreeding increases on smaller or more-visited islands, but this does not seem to be the case. Mice from Antipodes and Pitt Islands - both substantially smaller and more remote than Floreana - nonetheless have lower inbreeding coefficients than mice from Floreana (*p* < 0.01 in pairwise comparisons by Wilcoxon rank-sum tests, corrected for multiple testing). We note that although island land area and level of inbreeding have significantly nonzero rank-correlation (Spearman’s *ρ* = −0.354, *p* = 0.003), there is no linear relationship (Pearson’s *r* = −0.004, *p* = 0.97).

Inbreeding accelerates the erosion of genetic diversity by drift. The effect should be stronger on the X chromosome than the autosomes because the X is present in fewer copies in the population. On the two islands for which the most data is available, we find that a substantial proportion of the genome is fixed for a single haplotype. The population on Southeast Farallon Island has fixed 4.6% of the autosomes and 39.4% of the X chromosome; on Floreana Island, 6.3% of the autosomes and 45.2% of the X chromosome are fixed (**Figure S5** and **Table S4**). The longest fixed regions span 6.2 Mb on the autosomes and 13.0 Mb on the X chromosome. We cannot exclude the possibility that some of these regions were fixed by selection on locally-adaptive mutations rather than by drift alone.

### Linkage disequilibrium

Ascertainment bias in the content of the MUGA family of arrays complicates the interpretation of estimators of effective population size derived from the site frequency spectrum. However, patterns of linkage disequilibrium (LD) - which are somewhat more robust to SNP ascertainment - also carry information about effective population size, especially in the recent past. We calculated the decay of autosomal LD in island populations and representative mainland populations and used the equation of Sved (1971) to estimate effective population size. Consistent with previous work on wild populations of M. m. *domesticus* (Laurie et al. 2007), LD decays to background levels within approximately 60 kb in mainland populations (Greece, France, Maryland). (The median spacing of markers on our array is 32 kb.) LD is generally stronger in island populations, but the effect is most pronounced on Southeast Farallon and Floreana, in which LD persists over > 1 Mb (**Figure 5A**). Assuming a sex-averaged recombination rate of 0.52 cM/Mb (Liu et al. 2014), effective population sizes on these islands are only 273 ± 12 (mean ± SE) and 179 ± 8 individuals, respectively (**Figure 5B**). This is several orders of magnitude smaller than estimates for European *M. m. domesticus* based on either LD (this study) or neutral sequence (Geraldes et al. 2008), and approximately 400-fold smaller than the census size on Southeast Farallon at the peak of the breeding season.

These estimates of *N*_*e*_ from LD are based on a very simple demographic model of a single, closed population without selection. The reality is far more complicated, especially for the island populations of interest here. Migration may bring new haplotypes into the population, introducing an additional component of LD that is unrelated to the size of the existing population (*ie.* admixture LD). Population bottlenecks and directional changes in long-term population size over time also influence LD. To build qualitative intuition about the influence of these factors on LD in island mice, we performed coalescent simulations of a single ancestral population that splits into two (island and mainland) with a bottleneck on the island subpopulation, with varying degrees of migration after the split (**Figure S6**). LD is strongest - that is, decays most slowly over physical distance - in scenarios with a stronger colonization occurring more recently in the past. However, sufficient migration intensity (*Ne·m* ≥ 1) dramatically reduces the strength of LD. This implies that the effective population sizes on Southeast Farallon Island and Floreana Island are not only small but these populations have received few migrants since they were established.

**Figure 5:**
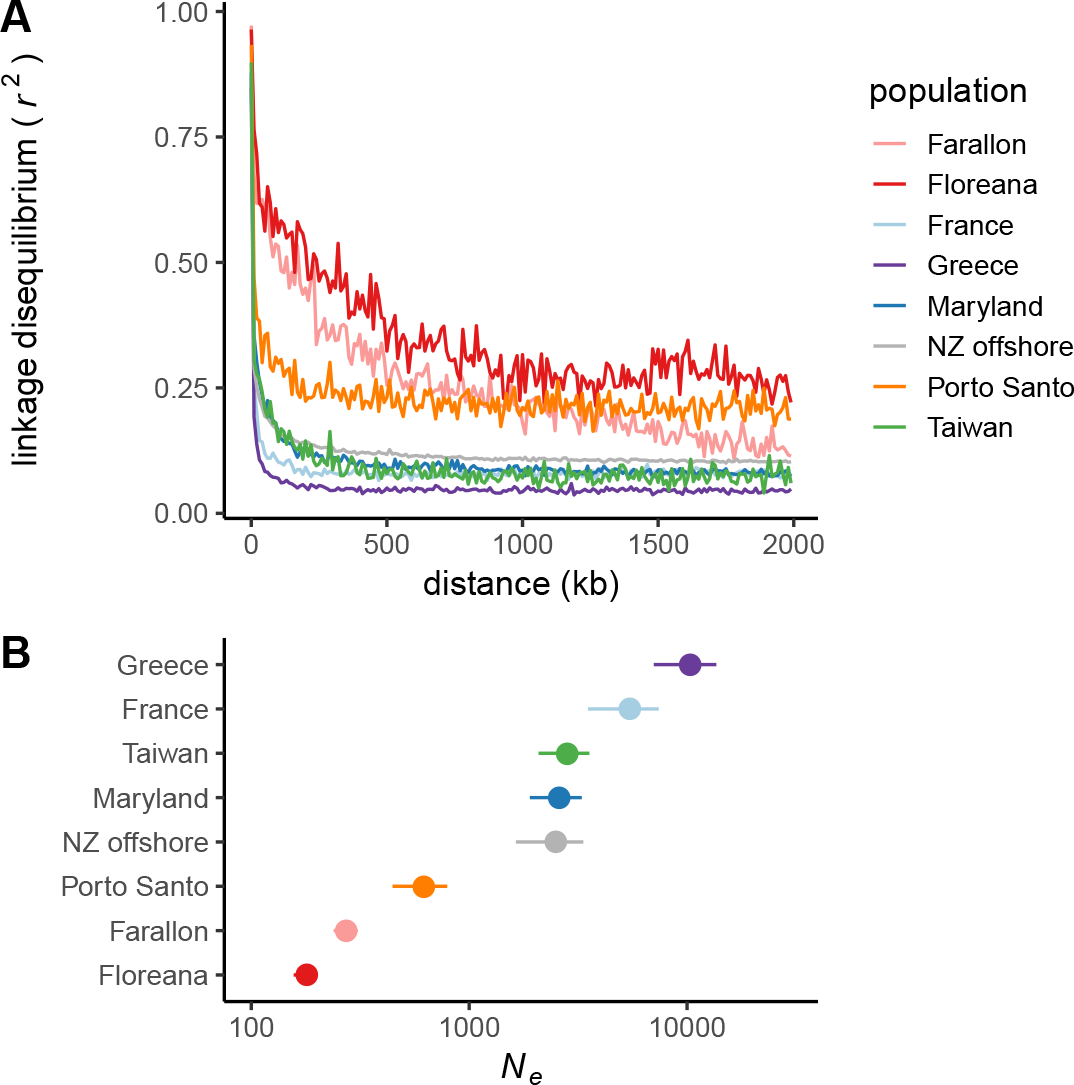
Decay of linkage disequilibrium **(A)** and corresponding estimates of effective population size **(B)** in island versus representative mainland populations.

### Rodenticide resistance

The most widely-used means of rodent eradication is application of coumarin-type anticoagulants such as warfarin and brodifacoum. These compounds irreversibly inhibit the vitamin K epoxide reductase complex (VKORC1) and cause death by hemorrhage in a dose-dependent manner (Rost et al. 2004). However, resistance alleles exist in multiple rodent species (Pelz et al. 2005). Our genotyping platform includes two SNPs in *Vkorcl* (p.Ala48Thr, p.Arg61Leu) that confer resistance to vitamin K antagonists. These alleles segregate at high frequency in *M. spretus* populations in Spain and northern Africa and have introgressed into *M. m. domesticus* under recent positive selection (Song et al. 2011). They are in strong LD in our dataset (*r*^2^ =0.89, *D’* = 1.0).

We find that the warfarin-resistant *M. spretus* haplotype tagged by these variants is common in Spain (derived allele frequency [DAF] = 0.50), Portugal (0.58) and France (0.16), rare in Maryland (0.03) and absent from all island populations studied (**Figure 6A**). The haplotype is likewise absent from *M. m. musculus* and *M. m. castaneus* in our cohort. This pattern is consistent with spatial diffusion of resistance alleles through Europe (**Figure 6B**).

**Figure 6:**
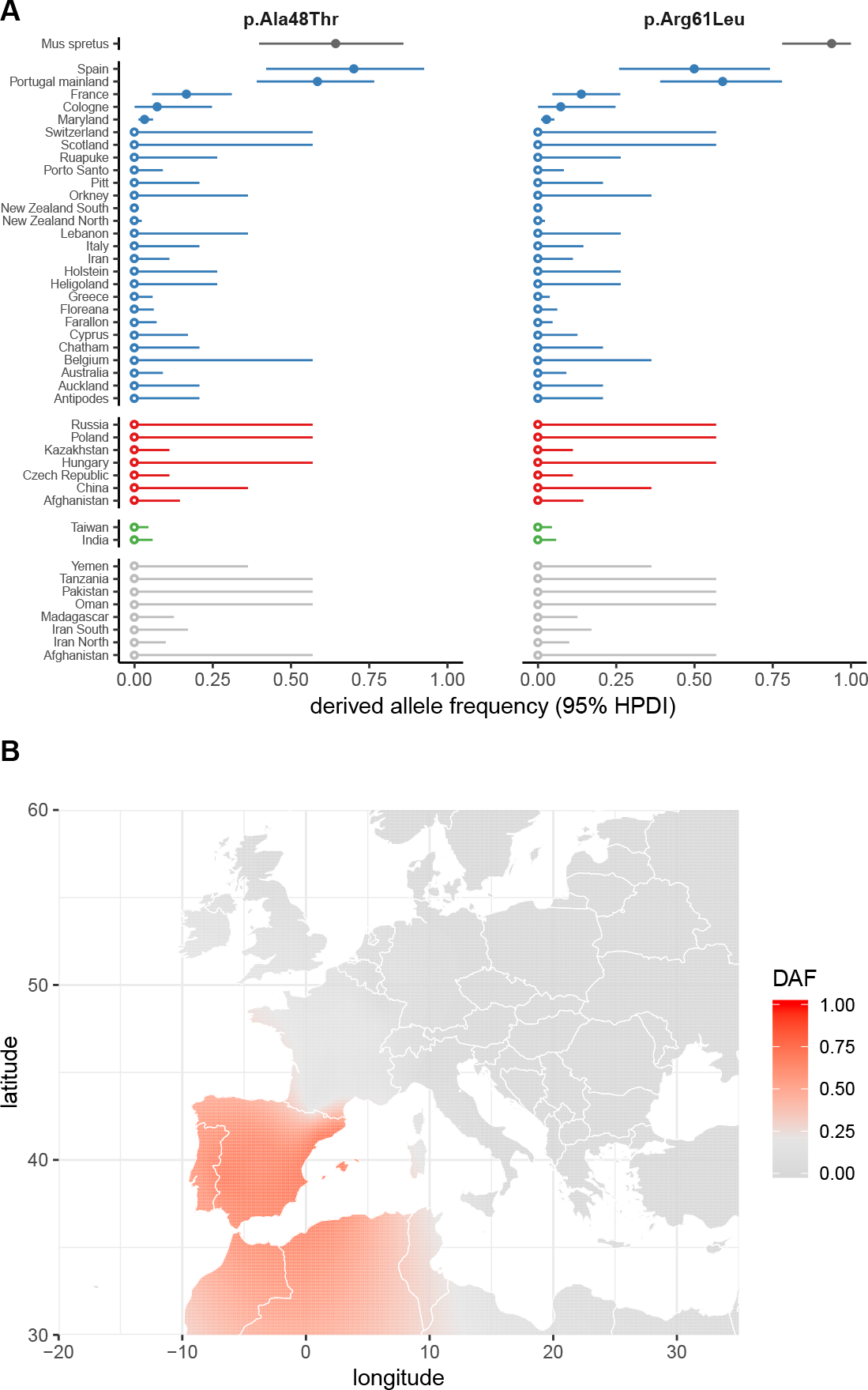
Distribution of anticoagulant resistance alleles. **(A)** Population-specific frequencies (with 95% highest posterior density intervals, HPDIs) of known *Vkorc1* alleles conferring resistance to coumarin-type anticoagulants including warfarin. **(B)** Spatially interpolated frequency of the p.Arg61Leu allele, demonstrating its spread outward from north Africa and southern Spain.

## Discussion

Here we present a detailed analysis of ancestry and population structure in house mice on offshore islands on the basis of genome-wide SNP data. Although the mitochondrial sequences used in previous studies of house mouse dispersal can reveal matrilineal origins in with great precision, the scope of inference possible from a single uniparentally-inherited locus is necessarily limited. Genome-wide SNP genotypes offer a richer and more nuanced portrait of ancestry that augments the historical record of dispersal events. Our characterization of the ancestry of house mice on offshore islands is nonetheless broadly consistent with an extensive literature on the dispersal of this species (Gabriel et al. 2010). Mice from all islands studied are of pure or very nearly pure *M. m. domesticus* origin (**Table S2**), irrespective of historical and ongoing contact with regions where *M. m. castaneus* or *M. m. musculus* predominate (Gabriel et al. 2010; King 2016). At the coarsest scale, two ancestry components - roughly corresponding to present-day Mediterranean and southern European populations, and to present-day populations in northern Europe, the British Isles and the Low Countries - are evident in multiple complementary analyses (**Figure 2, Figure 3**). This accords well with the synthesis of archaeological and mitochondrial evidence in favor of two waves of movement of house mice into Europe over the past 10,000 years, one by a southerly and one by a northerly route (Macholán et al. 2012). A more detailed analysis of population structure in Old World *M. m. domesticus* is provided in a companion manuscript (Hughes, Morgan, Didion *et al.*, forthcoming).

Individual-level ancestry profiles are consistent within geographically-defined populations on both the autosomes and X chromosome (**Figure S3**), supporting the notion that islands are resistant to repeated invasions after the initial colonization event (Hardouin et al. 2010; Jones et al. 2011; Gabriel et al. 2015). In a limited number of cases we can make a strong statement about the likely source population: Ruapuke appears to have been colonized from the eastern coast of Australia; Porto Santo from Portugal and Spain; and Heligoland from northern Germany (**Figure 3**). These findings are concordant with inferences from mitochondrial haplotypes. For the remaining islands, the picture is less clear - in part because genetic differentiation is limited between the source populations themselves, and in part because the ascertainment scheme for our array limits the number of population-specific alleles. Nonetheless we find that mice from islands with historical connections to the British Isles, such as Auckland, Chatham and Pitt in the Southern Ocean, have greatest affinity to present-day populations from northern Europe (Scotland, Belgium, Germany) and to larger landmasses colonized predominantly by the British (eastern United States, New Zealand) (**Figure 3**; see Veale et al. (2018) for more detailed treatment). Mice on Floreana Island, in the Galápagos, are clearly admixed, with substantial Iberian ancestry (**Figure 3**) but a northern European Y chromosome. The origins of Farallon mice are somewhat less clear. As for Floreana, models allowing for north-south admixture offer better fit the data than models without (**Figure S1**). But Farallon mice have less affinity to Iberian populations (**Figure 3**) in spite of carrying the “Mediterranean” Y haplogroup. An ancestry component that is prominent on Southeast Farallon does not appear in any of our mainland reference populations; this is a well-known property of STRUCTURE, ADMIXTURE and similar models in cases of strong population-specific drift (eg. after a bottleneck) (Puechmaille 2016). Understanding the origins of this component will ultimately require more thorough geographic sampling in Europe and the Americas. Finally, although we cannot exclude the possibility of multiple waves of invasion on either Farallon or Floreana, it seems more likely that admixture on both islands occurred prior to colonization - for instance in port cities where both food stores and migrants would be plentiful.

Inbreeding, as estimated from the proportion of the genome autozygous, arises both from non-random mating between close relatives and from endogamy in small populations. Values are high in all island populations surveyed but fall within the distribution of values for mainland populations (**Figure 4**). The level of inbreeding on islands has no apparent relationship to the land area, geographic isolation, latitude or other physical features of individual islands. This suggests that inbreeding on islands is driven by behavior or fine-scale features of the local landscape rather than broad geographic constraints. Indeed, it has been well-known since the earliest studies of protein allozymes (Philip 1938; Petras 1967) that wild house mouse populations are deficient in heterozygotes, consistent with local inbreeding. This accords well with the social structure of these populations: house mice live in small demes founded by up to a few dozen individuals that may cover as little as a 5 − 10 m^2^ (Berry 1970). Within these demes, males are fiercely territorial, and immigrants are rarely successful at joining in a new deme (Lidicker 1976). However, in well-connected landscapes, demes are sufficiently fluid over time and long-range dispersal (mostly by juvenile males (DeLong 1967; Pocock et al. 2005)) sufficiently frequent, that populations behaves as if they are approximately panmictic at the regional scale, despite being finely subdivided on the local scale (Nagylaki 1977; Berry 1986).

On isolated islands, the conditions underlying this approximation break down. Although demes may become genetically homogenized by within-island migration, heterozygosity decays if few migrants are exchanged with the mainland so that the effective population size remains small. We observe this most clearly on Southeast Farallon Island and Floreana Island (**Figure 5**). In this circumstance the level of genetic diversity is dominated by the strength of the bottleneck at the time the island was colonized, the time since colonization, and the size of the breeding population at its winter nadir. Because *N*_*e*_ is proportional to the harmonic mean of the number of breeding individuals across generations (Crow and Kimura 1970), which is in turn dominated by small values, seasonal booms in the census population size such as occur on Southeast Farallon have negligible impact on genetic diversity. (This can be confirmed by hand or by simulations; not shown.) On this basis we can conclude that both Southeast Farallon and Floreana were likely colonized by at most a few hundred breeding individuals each.

Our findings have implications for future eradication efforts. At this time the only feasible strategy for eradication is the use of rodenticide-laced bait, either by broadcast application or in feeding stations. This is usually effective when target animals are genetically susceptible (Howald et al. 2007), as we have shown in the case on Southeast Farallon and Floreana, but non-target species are also at risk (San Francisco Bay National Wildlife Refuge Complex 2013). Gene drives are being explored as a more targeted alternative (Backus and Gross 2016; Prowse et al. 2017; Leitschuh et al. 2018). One proposed technique is transgenic delivery of the male sex-determining factor (*Sry*) on in the background of the t-haplotype, an autosomal segregation distorter present at low frequencies in the wild (Backus and Gross 2016). This system would distort the sex ratio in favor of males and thus suppress the reproductive potential of the population, in similar manner to vector control systems recently trialled in mosquitos (Galizi et al. 2014). The efficiency of this approach would be blunted by the early loss of those alleles by drift at the small population sizes observed on Southeast Farallon and Floreana; on the other hand, small population size also allows low-fitness segregation distorters such as the *t^Sry^* construct to reach higher frequencies by drift (Lyon 2003), and mitigates the emergence of resistance (Unckless et al. 2017).

More broadly, our findings help to frame discussion of the “island syndrome” in murid rodents (Adler and Levins 1994). The most consistent and predictable feature of this syndrome in house mice is an allometric increase in body size (Pergams and Ashley 2001; Berry and Scriven 2005; Lomolino et al. 2013). Body size is the prototypical complex trait in mice and its genetics is relatively well-understood from both inbred line crosses and artificial selection on outbred stocks (among many references, Falconer (1973); Cheverud et al. (1996)). Crosses between inbred strains derived from Gough Island (GI) and a North American mainland population (WSB/EiJ) have provided empirical confirmation that the basis for island-mainland differences in body size - in at least one case - is polygenic, largely additive and shared between sexes (Gray et al. 2015). Such a rapid and vigorous response to selection in the face of low genetic variation post-bottleneck and inbreeding implies that selection must be very strong. But this interpretation is predicated on the assumption that larger body size confers higher fitness, which may or may not be the case for house mice (Ruff et al. 2017). Alternatively, the “island syndrome” may reflect a plastic response to the withdrawal of purifying selection and relaxation of sexual selection. How either scenario might interact with pervasive inbreeding (and inbreeding depression) - which we have shown to be the rule on islands - is an open question. Future studies would benefit from whole-genome sequencing to better characterize the genetic load and fine-scale patterns of variation in island populations.

## Acknowledgements

This work was supported by grants from the following federal agencies: DARPA: D16-59 (DWT); National Institutes of Health: F30MH103925 (APM), T32GM067553 (APM, JPD), U19AI100625 (FPMdV), P50GM076468 (Gary A Churchill); Oliver Smithies Investigator Award (FPMdV). We thank our many collaborators and sample collectors without whose extensive fieldwork this study would not have been possible, in particular the staff of Island Conservation on Southeast Farallon and Floreana. We thank Gerry McChesney of the US Fish and Wildlife Service for assistance with permits for trapping on Southeast Farallon Island. We thank the following technical staff for assistance with sample preparation, shipping and SNP array processing: Tim Bell, Ryan Buus, Jason Spence, Justin Gooch, Pablo Hock and Darla Miller. Finally, we are grateful to Andrew Veale for making genotypes from New Zealand populations available to the community, and for discussions regarding their history.

## Data accessibility

Genotypes will be deposited in Dryad in VCF and PLINK binary formats upon acceptance.

## Author contributions

Contributed to study design and sample collection: APM, JPD, JJH, JBS, WJJ, KJC, DWT, FPMdV. Analyzed data: APM, JPD, JJH. Wrote paper: APM, JPD.

## Supplementary material

**Table S1.** Full sample information. See included column key for details.

**Table S2.** Subspecies ancestry proportions from supervised ADMIXTURE analysis using a subset of 8 875 autosomal and 401 X-chromosomal subspecies-informative markers. One column each for point estimates and standard errors for *M. m. domesticus* (dom), *M. m. musculus* (mus) and *M. m castaneus* (cas) contribution. Nominal taxonomic designation of each individual based on geographic origin and morphology is also shown.

**Table S3.** Summary of mitochondrial and Y chromosome haplogroup assignments by taxon and population.

**Table S4.** Genomic regions fixed on Farallon and Floreana islands.

**File S1.** Genotype matrix including all individuals (*N* = 520).

**File S2.** Genotype matrix including only unrelated individuals (*N* = 403).

**Figure S1:**
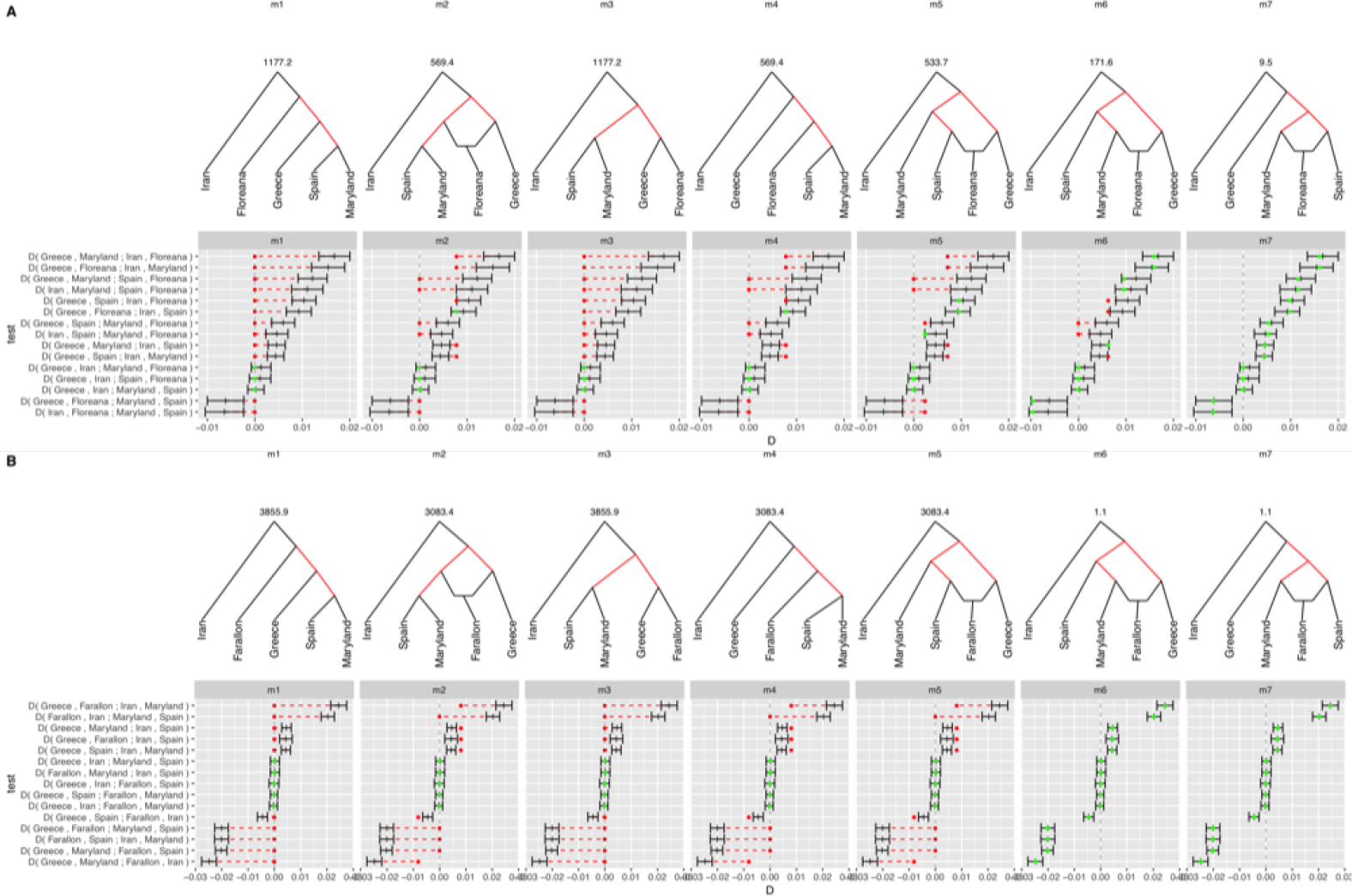
Model-based fitting of admixture graphs to *f*_4_-statistics (ie. *D*-statistics) calculated from observed allele frequencies. Admixture scenarios giving rise to the Floreana Island and Southeast Farallon Island populations are shown in panels **A** and **B**, respectively. In each panel, top row shows pre-specified topology of admixture graph; edges whose weight is estimable from the data are shown in red. Value of cost function for the fitted graph is shown above the graph. Bottom row of each panel shows expected values (round points) of *D*-statistics in the fitted graph compared to 95% confidence intervals about their observed values (vertical hashes). Expected values falling within the CI for the observed value are colored green; values falling outside the CI are colored red.

**Figure S2:**
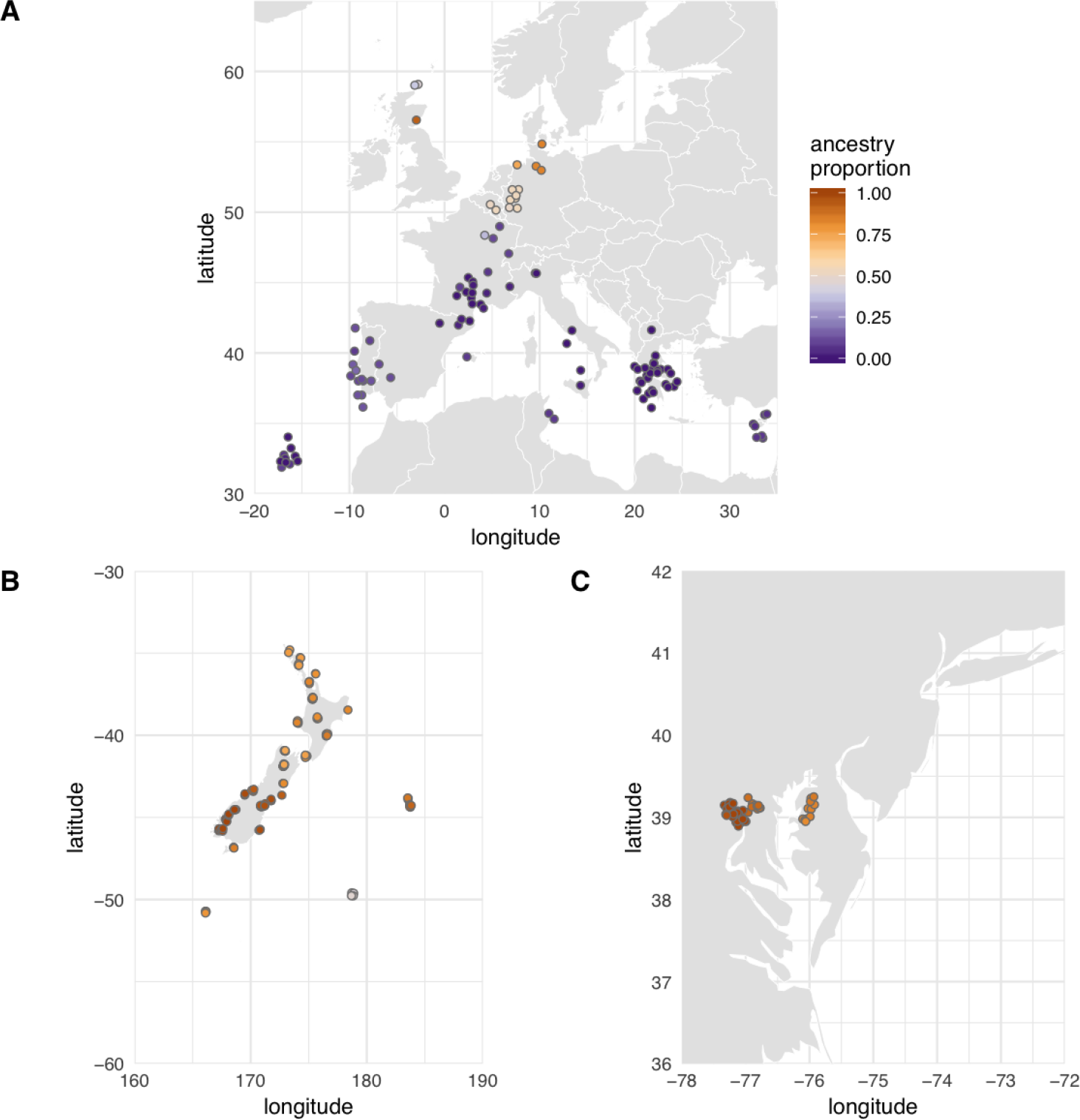
Proportion of “northern European”-like ancestry in ADMIXTURE analysis with *K* = 2 (pink bar in first row of **Figure S1**) for mice trapped in **(A)** Europe and the Mediterranean; **(B)** New Zealand and the Southern Ocean; and **(C)** Maryland, United States. A small amount of random “jitter” has been applied to each point to reduce overplotting, so locations displayed are not exact.

**Figure S3:**
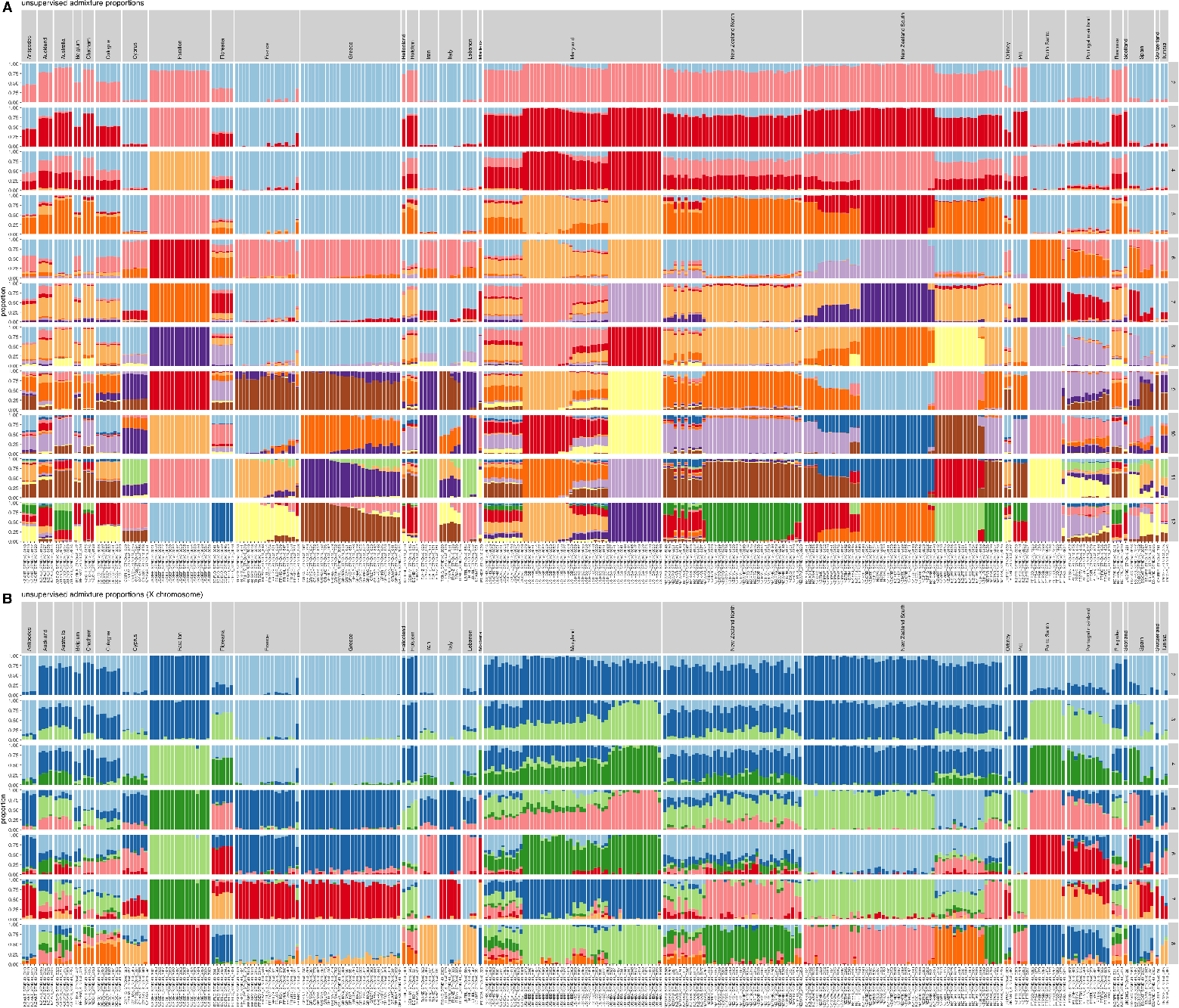
Barplots of ancestry proportions from unsupervised ADMIXTURE analyses of *M. m. domesticus* samples: **(A)** autosomes (*K* = 2,…, 12); **(B)** X chromosome (*K* = 2,…, 8). Individuals are shown in the same order in panels A and B.

**Figure S4:**
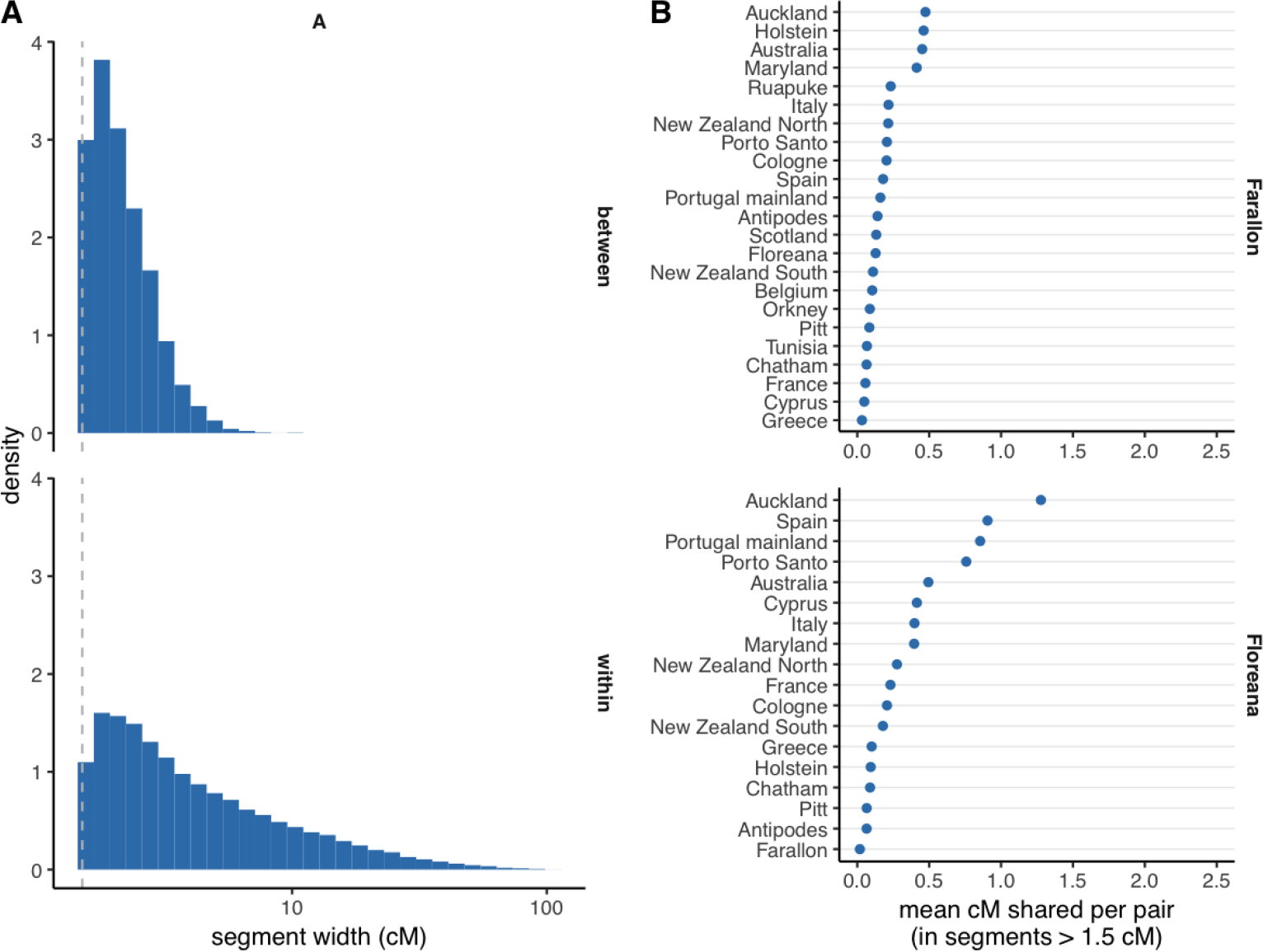
Sharing of long haplotypes identical-by-descent (IBD) within and between populations. **(A)** Distribution of size of autosomal segments shared IBD within (top) or between (bottom) populations. Dashed line marks 1.5 cM cutoff used for analysis described in main text. **(B)** Mean genomic territory covered by segments > 1.5 cM shared IBD between all pairs of mice from Farallon (top) or Floreana (bottom) with mice from populations shown on y-axis.

**Figure S5:**
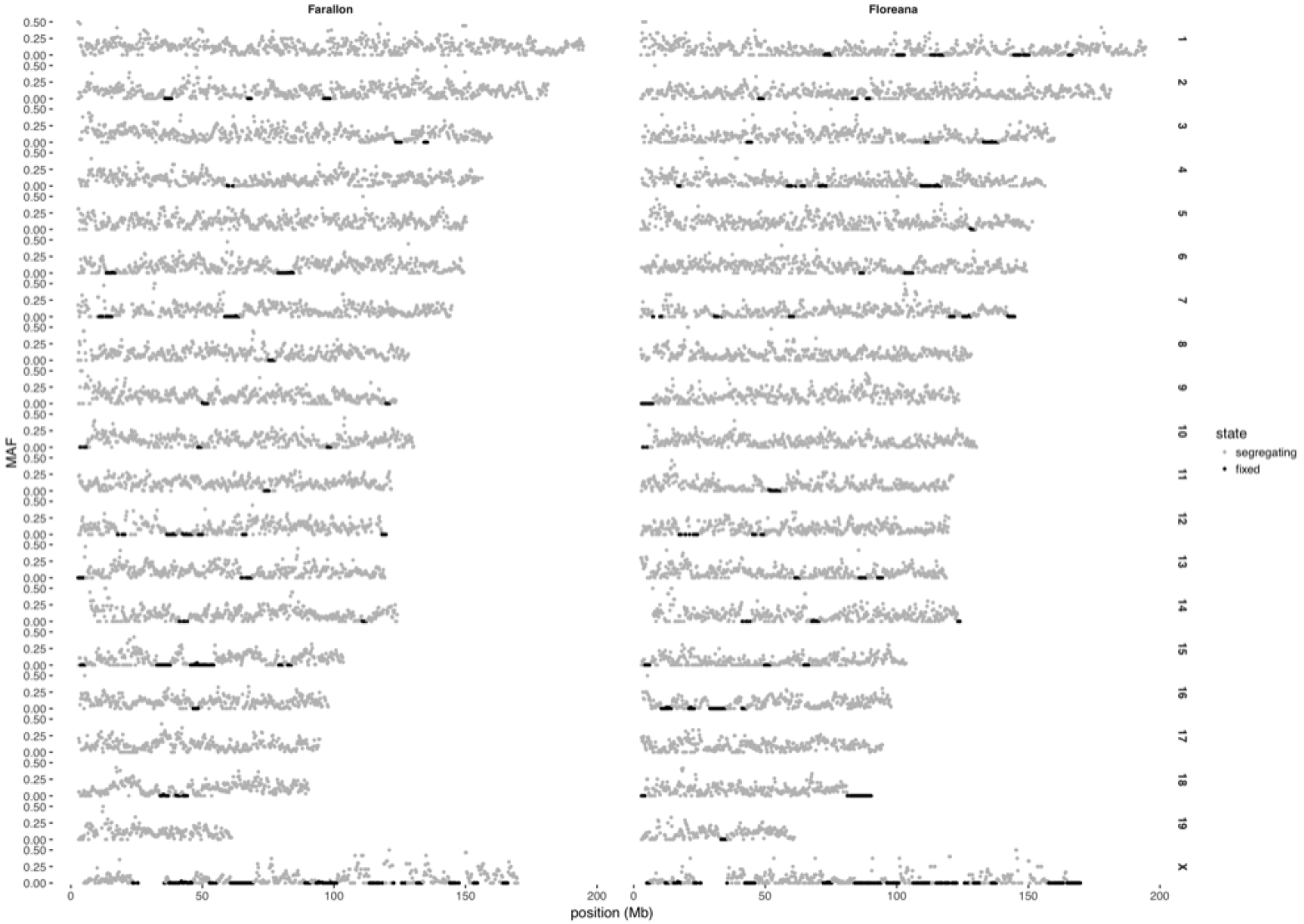
Regions fixed in Farallon or Floreana mice. Each point represents one 500 kb window; observed minor-allele frequency is plotted on the y-axis. Colors indicate binarized segregation status (segregating or fixed) inferred by HMM.

**Figure S6:**
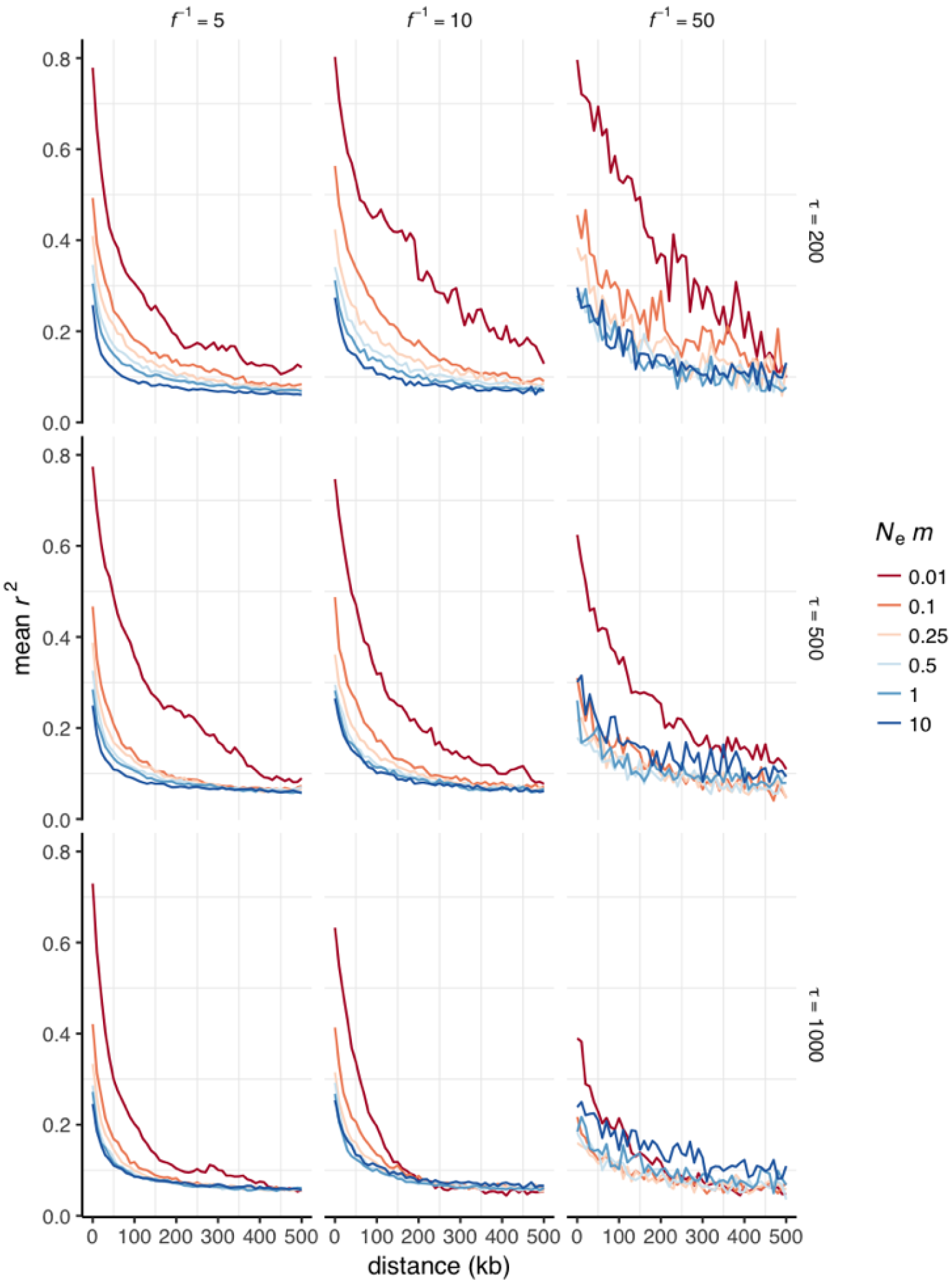
LD decay curves from coalescent simulations under island-mainland models. Each panel shows LD decay (according to effective migration rate *N*_*e*_·*m*) for a unique combination of strength of colonization bottleneck (*f*^−1^) and time since colonization (*τ*).

## References

Adler, GH, Levins, R. 1994. The island syndrome in rodent populations. Q. Rev. Biol. 69:473–490.

Alexander, DH, Novembre, J, Lange, K. 2009. Fast model-based estimation of ancestry in unrelated individuals. Genome Res. 19:1655–1664.

Arthur, R, Schulz-Trieglaff, O, Cox, AJ, O’Connell, J. 2017. AKT: ancestry and kinship toolkit. Bioinformatics. 33:142–144.

Backus, GA, Gross, K. 2016. Genetic engineering to eradicate invasive mice on islands: modeling the efficiency and ecological impacts. Ecosphere. 7.

Berry, RJ. 1970. The natural history of the house mouse. Field Studies. 3:219–262.

Berry, RJ. 1986. Genetical processes in wild mouse populations. past myth and present knowledge. In: The Wild Mouse in Immunology. Springer Berlin Heidelberg, pp. 86–94.

Berry, RJ, Scriven, PN. 2005. The house mouse: a model and motor for evolutionary understanding. Biol. J. Linn. Soc. Lond. 84:335–347.

Bishop, CE, Boursot, P, Baron, B, Bonhomme, F, Hatat, D. 1985. Most classical mus musculus domesticus laboratory mouse strains carry a mus musculus musculus Y chromosome. Nature. 315:70–72.

Browning BL, Browning, SR. 2013. Improving the accuracy and efficiency of Identity-by-Descent detection in population data. Genetics. 194:459–471.

Cheverud, JM, Routman, EJ, Duarte, FA, van Swinderen B, Cothran, K, Perel, C. 1996. Quantitative trait loci for murine growth. Genetics. 142:1305–1319.

Crow, JF, Kimura, M. 1970. An Introduction to Population Genetics Theory. New Jersey: The Blackburn Press.

DeLong, KT. 1967. Population ecology of feral house mice. Ecology. 48:611–634.

Didion JP, Morgan, AP, Yadgary, L, et al. (48 co-authors). 2016. R2d2 drives selfish sweeps in the house mouse. Mol. Biol. Evol. 33:1381–1395.

Didion, JP, Yang, H, Sheppard, K, Fu, CP, McMillan, L, de Villena FPM, Churchill, GA. 2012. Discovery of novel variants in genotyping arrays improves genotype retention and reduces ascertainment bias. BMC Genomics. 13:34.

Falconer, DS. 1973. Replicated selection for body weight in mice. Genet. Res. 22:291–321.

Gabriel, SI, Johannesdottir, F, Jones, EP, Searle, JB. 2010. Colonization, mouse-style. BMC Biol. 8:131.

Gabriel, SI, Mathias, ML, Searle, JB. 2015. Of mice and the ‘age of discovery': the complex history of colonization of the azorean archipelago by the house mouse (mus musculus) as revealed by mitochondrial DNA variation. J. Evol. Biol. 28:130–145.

Galizi, R, Doyle, LA, Menichelli, M, Bernardini, F, Deredec, A, Burt, A, Stoddard, BL, Windbichler, N, Crisanti, A. 2014. A synthetic sex ratio distortion system for the control of the human malaria mosquito. Nat. Commun. 5:3977.

Geraldes, A, Basset, P, Gibson, B, Smith, KL, Harr, B, Yu, HT, Bulatova, N, Ziv, Y, Nachman, MW. 2008. Inferring the history of speciation in house mice from autosomal, x-linked, y-linked and mitochondrial genes. Mol. Ecol. 17:5349–5363.

Gray, MM, Parmenter, MD, Hogan, CA, Ford, I, Cuthbert, RJ, Ryan, PG, Broman, KW, Payseur, BA. 2015. Genetics of rapid and extreme size evolution in island mice. Genetics. 201:213–228.

Gray, MM, Wegmann, D, Haasl, RJ, White, MA, Gabriel, SI, Searle, JB, Cuthbert, RJ, Ryan, PG, Payseur, BA. 2014. Demographic history of a recent invasion of house mice on the isolated island of gough. Mol. Ecol. 23:1923–1939.

Gündüz, I, Auffray, JC, Britton-Davidian, J, Catalan, J, Ganem, G, Ramalhinho, MG, Mathias, ML, Searle, JB. 2001. Molecular studies on the colonization of the madeiran archipelago by house mice. Mol. Ecol. 10:2023–2029.

Hardouin, EA, Chapuis, JL, Stevens, MI, van Vuuren JB, Quillfeldt, P, Scavetta, RJ, Teschke, M, Tautz, D. 2010. House mouse colonization patterns on the sub-antarctic kerguelen archipelago suggest singular primary invasions and resilience against re-invasion. BMC Evol. Biol. 10:325.

Harper, GA, Carrion, V. 2011. Introduced rodents in the Galápagos: colonisation, removal and the future. In: Veitch CR, Clout MN, Towns DR, editors, Island invasives: eradication and management, Gland, Switzerland: IUCN, pp. 63–66.

Harr, B, Karakoc, E, Neme, R, et al. (19 co-authors). 2016. Genomic resources for wild populations of the house mouse, mus musculus and its close relative mus spretus. Scientific Data. 3:160075.

Howald, G, Donlan, CJ, Galván, JP, et al. (12 co-authors). 2007. Invasive rodent eradication on islands. Conserv. Biol. 21:1258–1268.

Jones, EP, Jensen, JK, Magnussen, E, Gregersen, N, Hansen, HS, Searle, JB. 2011. A molecular characterization of the charismatic faroe house mouse. Biol. J. Linn. Soc. Lond. 102:471–482.

Kelleher, J, Etheridge, AM, McVean, G. 2016. Efficient coalescent simulation and genealogical analysis for large sample sizes. PLoS Comput. Biol. 12:e1004842.

King, CM. 2016. How genetics, history and geography limit potential explanations of invasions by house mice mus musculus in new zealand. Biol. Invasions. 18:1533–1550.

Laurie, CC, Nickerson, DA, Anderson, AD, Weir, BS, Livingston, RJ, Dean, MD, Smith, KL, Schadt, EE, Nachman, MW. 2007. Linkage disequilibrium in wild mice. PLoS Genet. 3:e144.

Leitschuh, CM, Kanavy, D, Backus, GA, Valdez, RX, Serr, M, Pitts, EA, Threadgill, D, Godwin, J. 2018. Developing gene drive technologies to eradicate invasive rodents from islands. Journal of Responsible Innovation. 5:S121–S138.

Leppälä K, Nielsen, SV, Mailund, T. 2017. admixturegraph: an R package for admixture graph manipulation and fitting. Bioinformatics. 33:1738–1740.

Lidicker, WZ. 1976. Social behaviour and density regulation in house mice living in large enclosures. J. Anim. Ecol. 45:677–697.

Liu, EY, Morgan, AP, Chesler, EJ, Wang, W, Churchill, GA, Villena, FPMd. 2014. High-Resolution Sex-Specific linkage maps of the mouse reveal polarized distribution of crossovers in male germline. Genetics. 197:91–106.

Lomolino, MV, van der Geer, AA, Lyras, GA, Palombo, MR, Sax, DF, Rozzi, R. 2013. Of mice and mammoths: generality and antiquity of the island rule. J. Biogeogr. 40:1427–1439.

Lyon, MF. 2003. Transmission ratio distortion in mice. Annu. Rev. Genet. 37:393–408.

Macholán, M, Baird, SJE, Munclinger, P, Piálek, J, editors. 2012. Evolution of the House Mouse. New York: Cambridge University Press, 1 edition edition.

Manichaikul, A, Mychaleckyj JC, Rich, SS, Daly, K, Sale, M, Chen, WM. 2010. Robust relationship inference in genome-wide association studies. Bioinformatics. 26:2867–2873.

McKenna, A, Hanna, M, Banks, E, et al. (11 co-authors). 2010. The genome analysis toolkit: a MapReduce framework for analyzing next-generation DNA sequencing data. Genome Res. 20:1297–1303.

McQuillan, R, Leutenegger, AL, Abdel-Rahman, R, et al. (21 co-authors). 2008. Runs of homozygosity in european populations. Am. J. Hum. Genet. 83:359–372.

Mills, K. 2016. Seabirds as part of migratory owl diet on southeast farallon island, california. Mar. Ornithol. 44:121–126.

Morgan, AP. 2016. argyle: An R package for analysis of illumina genotyping arrays. G3. 6:281–286.

Morgan, AP, Fu, CP, Kao, CY, et al. (23 co-authors). 2016. The mouse universal genotyping array: From substrains to subspecies. G3. 6:263–279.

Nagylaki, T. 1977. Decay of genetic variability in geographically structured populations. Proc. Natl. Acad. Sci. U. S. A. 74:2523–2525.

Narasimhan, V, Danecek, P, Scally, A, Xue, Y, Tyler-Smith, C, Durbin, R. 2016. BCFtools/RoH: a hidden markov model approach for detecting autozygosity from next-generation sequencing data. Bioinformatics. 32:1749–1751.

Neme, R, Tautz, D. 2016. Fast turnover of genome transcription across evolutionary time exposes entire noncoding DNA to de novo gene emergence. Elife. 5:e09977.

Pebesma, EJ. 2004. Multivariable geostatistics in s: the gstat package. Comput. Geosci. 30:683–691.

Pelz, HJ, Rost, S, Hünerberg, M, et al. (11 co-authors). 2005. The genetic basis of resistance to anticoagulants in rodents. Genetics. 170:1839–1847.

Pergams, OR, Ashley, MV. 2001. Microevolution in island rodents. Genetica. 112-113:245–256.

Peter, BM. 2016. Admixture, population structure, and F-Statistics. Genetics. 202:1485–1501.

Petras, ML. 1967. STUDIES OF NATURAL POPULATIONS OF MUS. i. BIOCHEMICAL POLYMORPHISMS AND THEIR BEARING ON BREEDING STRUCTURE. Evolution. 21:259–274.

Philip, U. 1938. Mating systems in wild populations of dermestes vulpinus and mus musculus. J. Genet. 36:197–211.

Pickrell, JK, Pritchard, JK. 2012. Inference of population splits and mixtures from Genome-Wide allele frequency data. PLoS Genet. 8:e1002967.

Pocock, MJO, Hauffe, HC, Searle, JB. 2005. Dispersal in house mice. Biol. J. Linn. Soc. Lond. 84:565–583.

Prowse, TAA, Cassey, P, Ross, JV, Pfitzner, C, Wittmann, TA, Thomas, P. 2017. Dodging silver bullets: good CRISPR gene-drive design is critical for eradicating exotic vertebrates. Proc. Biol. Sci. 284.

Puechmaille, SJ. 2016. The program structure does not reliably recover the correct population structure when sampling is uneven: subsampling and new estimators alleviate the problem. Mol. Ecol. Resour. 16:608–627.

Rost, S, Fregin, A, Ivaskevicius, V, et al. (13 co-authors). 2004. Mutations in VKORC1 cause warfarin resistance and multiple coagulation factor deficiency type 2. Nature. 427:537–541.

Ruff, JS, Cornwall, DH, Morrison, LC, Cauceglia, JW, Nelson, AC, Gaukler, SM, Meagher, S, Carroll, LS, Potts, WK. 2017. Sexual selection constrains the body mass of male but not female mice. Ecol. Evol. 7:1271–1275.

San Francisco Bay National Wildlife Refuge Complex. 2013. South farallon islands invasive house mouse eradication project: Revised draft environmental impact statement. Technical Report 78 FR 50082, US Fish and Wildlife Service.

Searle, JB, Jamieson, PM, Günduz, I, Stevens, MI, Jones, EP, Gemmill, CEC, King, CM. 2009. The diverse origins of new zealand house mice. Proc. Biol. Sci. 276:209–217.

Song, Y, Endepols, S, Klemann, N, Richter, D, Matuschka, FR, Shih, CH, Nachman, MW, Kohn, MH. 2011. Adaptive introgression of anticoagulant rodent poison resistance by hybridization between old world mice. Curr. Biol. 21:1296–1301.

Steiner, J. 1989. Temporal and spatial distribution of the camel cricket, farallonophilus cavernicola rentz (orthoptera: Gryllacrididae), on southeast farallon island, california. Pan-Pac Entomol. 65.

Sved, JA. 1971. Linkage disequilibrium and homozygosity of chromosome segments in finite populations. Theor. Popul. Biol. 2:125–141.

Thompson, BYCH. 1896. EGG-HUNTING ON THE SOUTH FARALLON. Frank Leslie’s Popular Monthly (1876-1904); New York. XLII:0_030.

Unckless, RL, Clark, AG, Messer, PW. 2017. Evolution of resistance against CRISPR/Cas9 gene drive. Genetics. 205:827–841.

Veale, AJ, Russell, JC, King, CM. 2018. The genomic ancestry, landscape genetics and invasion history of introduced mice in new zealand. R Soc Open Sci. 5:170879.

White, P. 1995. The Farallon Islands: Sentinels of the Golden Gate. Scottwall Associates.

